# Dissociable neurofunctional and molecular characterizations of reward and punishment sensitivity

**DOI:** 10.1101/2024.12.30.630747

**Authors:** Ting Xu, Chunhong Zhu, Xinqi Zhou, Zhiyi Chen, Xianyang Gan, Xiaobing Cui, Feng Zhou, Ran Zhang, Weihua Zhao, Xiaodong Zhang, Hong Chen, Qinghua He, Xu Lei, Jiang Qiu, Tingyong Feng

## Abstract

While the hyper-and hypo-reward or punishment sensitivities (RS, PS) have received considerable attention as prominent transdiagnostic features of psychopathology, the lack of an overarching neurobiological characterization currently limits their early identifications and neuromodulations. Here we combined microarray data from the Allen Human Brain Atlas with a multimodal fMRI approach to uncover the neurobiological signatures of RS and PS in a discovery-replication design (N=655 participants). Both RS and PS were mapped separately in the brain, with the intrinsic functional connectome in the fronto-striatal network encoding reward responsiveness, while the fronto-insular system was particularly engaged in punishment sensitivity. This dissociable functional connectome patterns related to RS and PS were also specific in differentiating decisions driven by social or monetary reward and punishment motivations. Further imaging transcriptomic analyses revealed that functional connectome variations for RS and PS were associated with topography of specific gene sets enriched in ontological pathways, including synaptic transmission, dopaminergic metabolism, immune response and stress adaptation. On the neurotransmitter level, the serotonin neuromodulator was identified as a pivotal hub regulating the intrinsic functional connectome patterns of RS and PS, with this process critically dependent on its interactions with dopaminergic, opioid and GABAergic systems. Overall, these findings indicate dissociable neural connectome mapping of RS and PS and highlight their linkage with transcriptomic profiles, which may offer valuable insights into the treatment evaluation for symptomatology relevant to reward/punishment processing deficits.

## INTRODUCTION

Selecting actions that maximize rewards and minimize punishments or losses within a computational framework of reinforcement learning is a hallmark of biological agents, which allows them to effectively interact with complex environments and develop adaptive behaviors ^1^. Achieving such goals specifically relies on the degree to which an individual’s behavior is motivated by rewards (i.e., sensitivity to reward, RS) or inhibited by the punishments (sensitivity to punishment, PS). However the functional roles of RS and PS are variable because individuals take different considerations on reward and punishments ^2,3^. For instance, traditional value-based learning models propose that individuals act as safety-seekers, assigning greater weight to potential losses than gains, even when the expected values of the outcomes are identical ^4,5^. This asymmetry in the evaluation of rewards and punishments is also evident when individuals depend on reward and punishment feedbacks to guide their choices. Specifically the increases in reward magnitude can promote repeated rewarding choices, whereas punishment avoidance remains unaffected by changes in penalty magnitude ^6^. Within those contexts, normal engagements of RS and PS allow beneficial decisions, such as reducing high-risk investment strategies and increasing the pursuit of rewards. However the abnormal changes of RS and PS may serve as candidate mechanisms for motivational and emotional dysregulations in some psychiatric disorders. Accumulate clinical research has highlighted the central role of reward hyposensitivity in both anhedonia and the overall depressive symptomatology ^7,8^, whereas suggested that heightened sensitivity to punishments exacerbated symptoms of rumination and worry in depression and anxiety disorders ^9,10^. However, an integrative and overarching biological-informed comprehension about the RS and PS is currently lacking, while clarifying it can promote the progress of early identification and intervention research.

As fundamental principles that govern behavioral adaptation to the environment, RS and PS are mediated by dissociable yet interconnected neural systems ^8,11^. Different lines of animal models combining with electrophysiological approaches have supported anatomical dissociation between RS and PS by revealing engagement of anatomically segregated midbrain neurons in processing reward and aversive stimuli ^9,12^. Consistent with this, remarkable observations in human fMRI works also demonstrate preferential activation of ventral parts of frontostriatal circuits during reward seeking behaviors, whereas concurrently implicating the dorsal regions in punishment avoidance ^13,14^. Furthermore individual differences in RS were also positively associated with activation in the ventral striatum during reward reception, while the individual differences in PS showed a negative association with activation in the left dorsal striatum during loss avoidance anticipation ^2^. However there is also evidence that punishment avoidance involves regions outside the frontostriatal projections of the brain-stem neuromodulator systems. According to this view, predisposition to detect and process punishments is additionally modulated by aversive signals encoded in other cortical and subcortical areas, such as the orbitofrontal cortex (OFC) and insula ^15,16^. The insula, in particular, has been shown to represent cues predicting primary punishments, such as disgust tastes and smells ^17,18^, with these findings later extending to more abstract types of punishments, including violations of social-fairness norms and monetary loss ^16,19,20^. In contrast to the role of insula in directly signaling aversive events via possible processes of salience detection ^21,22^ and interoception ^23,24^, the neurons of OFC are more sensitive to the outcome values ^25^ and are involved in the cognition evaluation of punishment through joint integration with other prefrontal regions ^26^.

For example, both the OFC and dorsolateral prefrontal cortex are crucial for recognizing violations of expected values^27^, with increased activation occurring in response to the absence of an expected reward (loss) or the presence of an unexpected electric shock (punishment)^28–30^. While these works have underscored the pivotal contribution of fronto-striatal and fronto-insular circuitry to regulate RS and PS, the underlying biological mechanisms behind the modulation of those neural systems to RS and PS changes remain under-investigated.

In fact, a plethora of animal and human works have outlined both RS and PS as evolutionary important constructs which can be heritable and are sub-served by relatively distinct transcriptional architectures ^31,32^. There is compelling evidence that both dopaminergic and opioid systems contribute to the experience of reward and motivation but in dissociable and complementary ways ^33–35^. Specifically, dopamine is primarily involved in the motivational aspects of reward, driving approach behaviors to attain the desired goal ^36,37^, whereas once the reward is obtained, the opioid system enhances the emotional and hedonic experience of that reward, reinforcing the behavior that led to it ^38^. In addition to dopamine and opioid system, recent pharmacological studies bring a new line of discovery on the key role serotonin plays in reward processing ^39,40^. By utilizing optogenetic manipulation of genetically identified cells, researchers have demonstrated that activating serotonergic neurons stimulates dopaminergic neurons via glutamatergic co-transmission and 5-HT receptor activation, thereby leading to reward ^41^. Serotonin is also strongly implicated in punishment ^42^, with proposed effects in enhancing instrumental learning from monetary loss ^43^ and facilitating responses to aversive stimuli ^44,45^. Lesions of serotonin containing terminals, systemic injections 5-HT antagonists or 5-HT synthesis inhibitors all exhibited anti-punishment effects ^46,47^, while a dietary manipulation by acute tryptophan depletion putatively impairing 5-HT transmission also reduced the suppression of aversively-motivated behavior ^48^. Additionally the interaction of serotonin system with prefrontal GABAergic neurotransmission also affected the processing of aversive and threatening environmental stimuli ^49^. However how those neuromodulatory transmitters are spatially distributed in the brain to guide individual responsiveness to reward and punishments has not been examined.

Recent advances in imaging transcriptomic approaches allow to specify the underlying biological pathways of macroscopic neuroimaging phenotypes ^50^, which may help elucidate how molecular transcriptomic variations impart brain functions relevant to RS and PS. By leveraging this sophisticated method in which the multivariate partial least squares regression (PLSR) was used to link the spatial expression of genes from Allen human brain atlas (AHBA) ^51^ with brain structural or functional patterns, researchers can indicate how alterations at the microscale architecture drive macroscale brain properties changes. This methodology has the potential to bridge the gap in identifying plausible transcriptomic correlates of neuroimaging biomarkers ^50^, holding promise to: 1) enhance understanding of genetic contributions to regional vulnerability in brain diseases ^52–55^, and 2) offer comprehensive biological targets for drug development and translation ^56,57^. In this context, several supportive works have uncovered that neurofunctional alterations related to reward motivation deficits in depression linked with regional expression of dopamine and 5HT neurotransmitters ^58,59^. Anatomical distribution of novel neuropeptides signaling such as oxytocin and angiotensin II type 1 receptor has been characterized in the brain and were specifically overlapped with functional imaging maps categorized as anticipation and learning of reward and aversive events ^56,57^.

In this study, we included two independent parts of an extensive resting state fMRI datasets as discovery (N=427 healthy participants) and replication (N=228 healthy participants) samples, and combined with the AHBA microarray data to identify distinct neurobiological signatures of RS and PS via multimodal analyses on brain functional connectome, genetic and transcriptional levels. Specifically, we employed the intersubject representational similarity analysis (IS-RSA) that allows the identification of shared neural representations for a given cognitive process while considering the individual variation simultaneously ^60,61^, to examine which brain regions display similar resting-state functional connectome (rs-FC) patterns in response to RS and PS, respectively (**Fig. 1a**). Those functional connectome signatures of RS and PS were further applied in two independent task fMRI datasets (S1: N=52, S2: N=37, S3: 6 participants from S1 were excluded due to imaging data loss and only N=46 healthy participants were included, **Fig. 1b**) to validate their specificity within the contexts of social and monetary reward or punishments. Next, we utilized the PLSR to determine the anatomical expression of which specific genes can predict RS-or RS-related rs-FC variation, and those gene sets were further included into functional enrichment analyses to indicate the potential biological pathways (**Fig. 1c**). Finally in order to decode the distinct rs-FC patterns of RS and PS by chemoarchitectures, we investigated the relationship between neurotransmitter systems and rs-FC patterns of RS or PS (**Fig. 1c**).

**Fig. 1.**
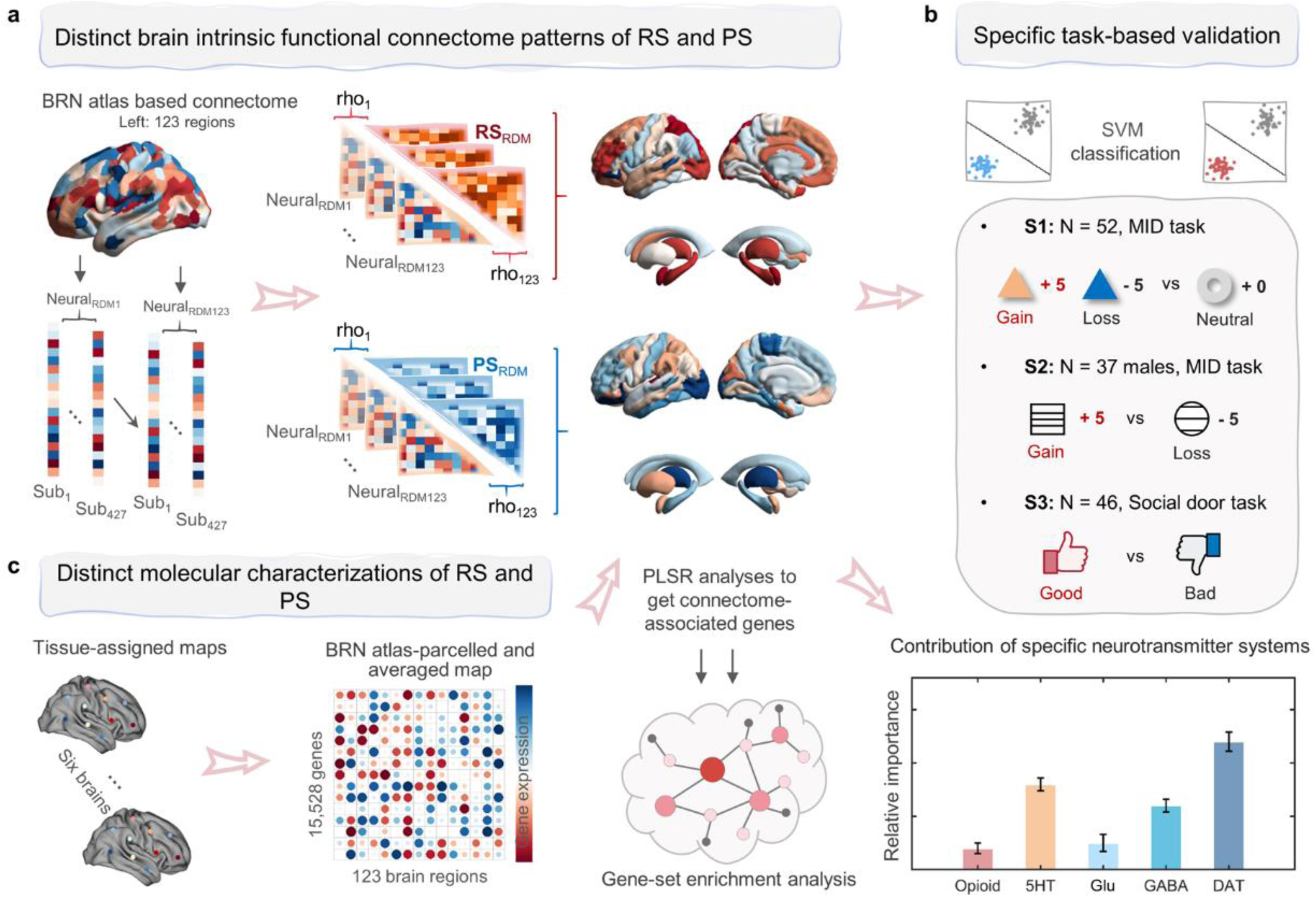
Schematic depiction of the analytic workflow. This study aims to: **(a)** identify the dissociable resting state brain functional connectome signatures of reward and punishment sensitivities via the use of inter-subject representational similarity analysis, **(b)** validate the specificity of brain functional connectome signatures of reward and punishment sensitivities in the contexts of social and monetary reward and punishment, respectively, **(c)** determine the anatomical expression of which specific genes from Allen human brain atlas can predict reward-or punishment sensitivities-related functional connectome variation, and those gene sets were further included into functional enrichment analyses to indicate the potential biological pathways, and finally examine the relative contribution of specific neurotransmitter systems to predict the brain functional connectome variation of reward and punishment sensitivities. Abbreviations: BRN-BrainNetome atlas, RDM-Representational dissimilarity matrix, SVM-Support vector machine, MID-Monetary incentive delay

## RESULTS

### Brain functional connectome signatures of RS and PS

Individual differences in sensitivity to reward and punishment have been extensively documented in past several years ^62^, with growing evidence highlighting the role of separable sets of neural regions in shaping these variations (e.g., activity of ventral striatum positively predicts RS while increased activation of the insula links with lower PS ^15,63^). However the traditional massive univariate ways that averaging the activity or connectivity strength across participants to some extent obscure the meaningful individual variability in how people process the same stimuli. To bridge this gap, we employed the intersubject representational similarity analysis which allows the identification of shared neural representations for a given cognitive process while considering the individual variation simultaneously ^60,61,64^, to examine which brain regions display similar rs-FC patterns in response to intersubject variability of RS or PS. Our results revealed that intersubject variation in RS score was positively reflected in brain regions within the fronto-striatal circuitry, including subcortical regions such as the thalamus, caudate and putamen, as well as frontal regions such as anterior cingulate cortex, ventromedial prefrontal cortex (vmPFC) and dorsomedial prefrontal cortex (All PS<0.05, Benjamini–Hochberg correction, **Fig. 2a**). In addition, several key regions within the frontal-insular circuitry encompassing inferior frontal gyrus, dlPFC, middle cingulate cortex and insula exhibited high representation similarity for the intersubject variation in PS score (All PS<0.05, Benjamini– Hochberg correction, **Fig. 2a**). Overall those results indicate the dissociable brain functional connectome signature of RS and PS, as evidenced by that the rs-FC patterns within fronto-striatal network encodes reward responsiveness while the fronto-insular system is more involved in the sensitivity to punishment.

**Fig. 2.**
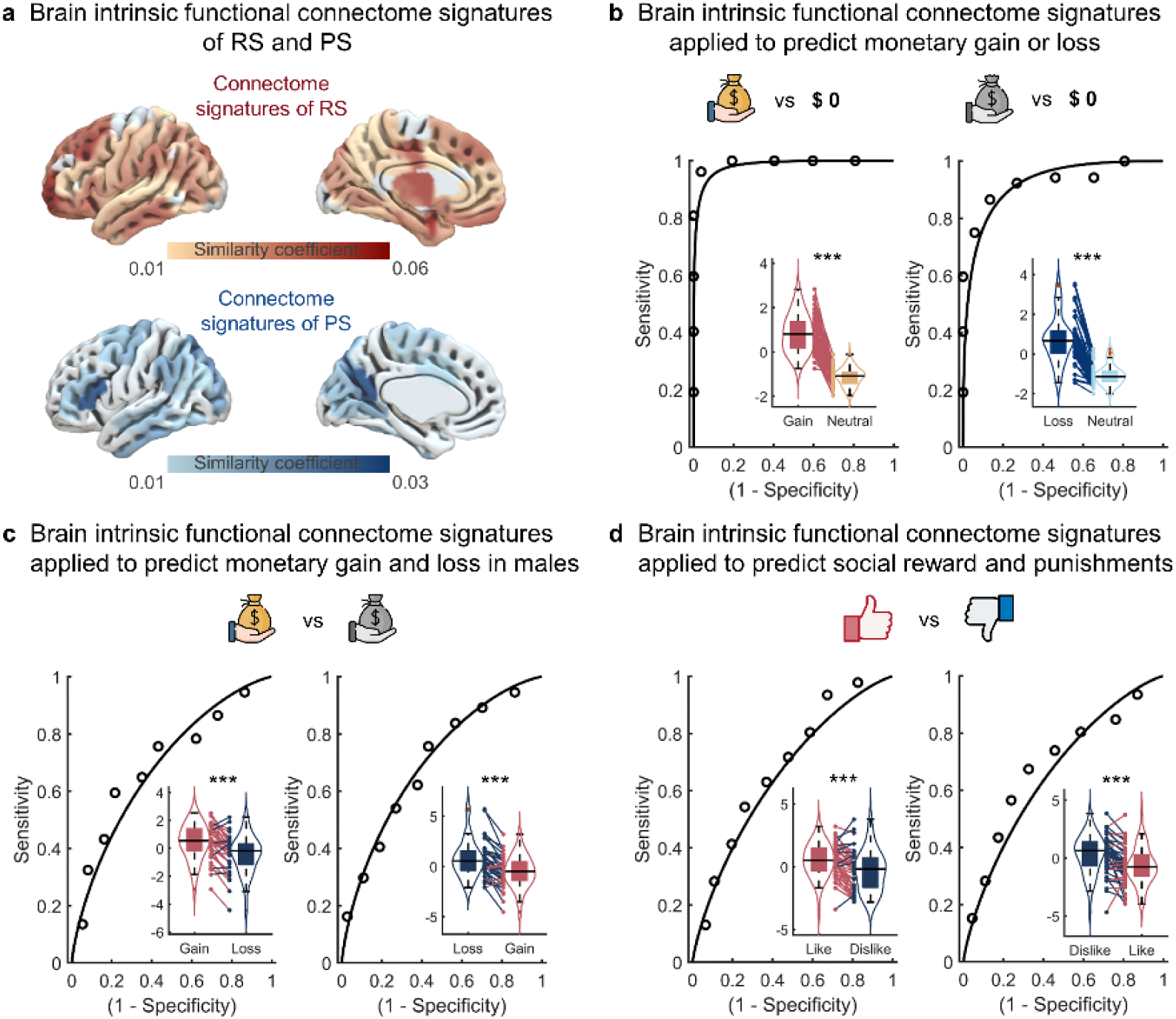
Brain functional connectome signatures of reward and punishment sensitivities, and its specificity within social/monetary reward and punishment contexts. (a) Regional functional connectome maps for reward sensitivity and punishment sensitivity are spatially distinct. **(b-d)** Robust specificity of the brain functional connectome signatures of reward and punishment sensitivities in the contexts of social and monetary reward or punishments, separately. The violin and box plots show the distributions of multivariate neural pattern map response to classify multiple events (i.e., Monetary gain vs neutral, Monetary loss vs neutral; Monetary gain vs loss, Monetary loss vs gain; Social like vs dislike, Social dislike vs like). The box is bounded by the first and third quartiles, and the whiskers stretched to the greatest and lowest values within the median±1.5 interquartile range, while each colored line between dots represents each participant’s paired. ***p<0.001

### Specificity of brain functional connectome signatures of RS and PS in monetary or social reward and punishment contexts

While the separate brain functional connectome signatures of RS and PS were primarily identified, we further applied those two signatures in the independent fMRI datasets (S1: N=52) that acquired neural responses during evaluation of monetary gain, loss and neutral (no gain or no loss) feedbacks, to explore their specificity within the contexts of monetary reward or punishments. As shown in **Fig. 2b**, the brain functional connectome signatures of RS can predict gain versus neutral events with high accuracy (accuracy=0.97, P<0.001; binominal test, t=21.22, p<0.001, Cohen’s d=1.65), whereas the brain functional connectome signatures of PS can significantly differentiate loss versus neutral events (accuracy=0.88, P<0.001; binominal test, t=15.68, p<0.001, Cohen’s d=1.42). To further test whether the prediction of brain functional connectome signatures of RS and PS in capturing reward or punishments can be segregate in monetary and social situations, we applied the connectome signatures of RS and PS to another two independent fMRI datasets acquired while participants directly received monetary gain or loss outcomes (S2: N=37), and social like or dislike labels (S3: 6 participants were excluded from S1 due to lack of task imaging data, leaving a final sample size of N=46). Our results showed that the brain functional connectome signatures of RS successfully differentiated the rewarding feedbacks from the punishment feedbacks regardless of monetary (accuracy=0.69, P=0.001; binominal test, t=6.62, p<0.001, Cohen’s d=0.66, **Fig. 2c**) or social situations (accuracy=0.66, P=0.007; binominal test, t=7.15, p<0.001, Cohen’s d=0.71, **Fig. 2c**). Similarly, the brain functional connectome signatures of PS were also able to distinguish monetary (accuracy=0.64, P=0.008; binominal test, t=5.62, p<0.001, Cohen’s d=0.56, **Fig. 2d**) or social punishments (accuracy=0.68, P<0.001; binominal test, t=5.32, p<0.001, Cohen’s d=0.57, **Fig. 2d**) versus corresponding rewards. In summary, these findings support the notion that the brain functional connectome signatures of RS and PS are not only robust in differentiating specific feedback types within the domains of monetary decision-making, but also generalized across social reward and punishment contexts.

### Regional gene expression linked with RS- and PS-related brain functional connectome alterations

Previous animal works indicate that the complex genetic variations influencing activity of specific neural ensembles is the underlying biological mechanism of hyper-or hypo-sensitivity to reward and punishments ^65–67^, yet the specific molecular underpinnings that shape distinct neural functional representations of RS and PS in humans remain elusive. In this regard, we utilized a whole brain transcriptomic dataset (i.e., AHBA) to characterize the gene expression patterns in the brain and further employed the PLSR for identifying the expression patterns of which specific gene sets were associated with the brain intrinsic functional connectome alterations of RS and PS, respectively. Specifically, by ranking all genes based on their multivariate correlations with the connectome pattern distributions of RS, we found a ranked gene list – PLS1– that explained 21.37% variance of the brain intrinsic functional connectome changes of RS. The spatial distribution of PLS1 was further positively correlated with the brain intrinsic functional connectome changes of RS (r_(121)_=0.33, P_spin_=0.01, corrected for multiple comparisons by spatial permutation testing (spin test)^68,69^, **Fig. 3a**) which means that genes positively (negatively) weighted on PLS1 are densely expressed in regions exhibiting highly similar (dissimilar) rs-FC patterns to represent RS. While applying the FDR correction to the normalized weights of PLS1, 204 positive weighted genes (Z>3.01) and 396 negative weighted genes (Z<-3.01, P_FDR_<0.005) were remained. We finally capitalized on the Metascape to perform gene set enrichment analysis for providing the biological and functional pathways of the PLS1.

**Fig. 3.**
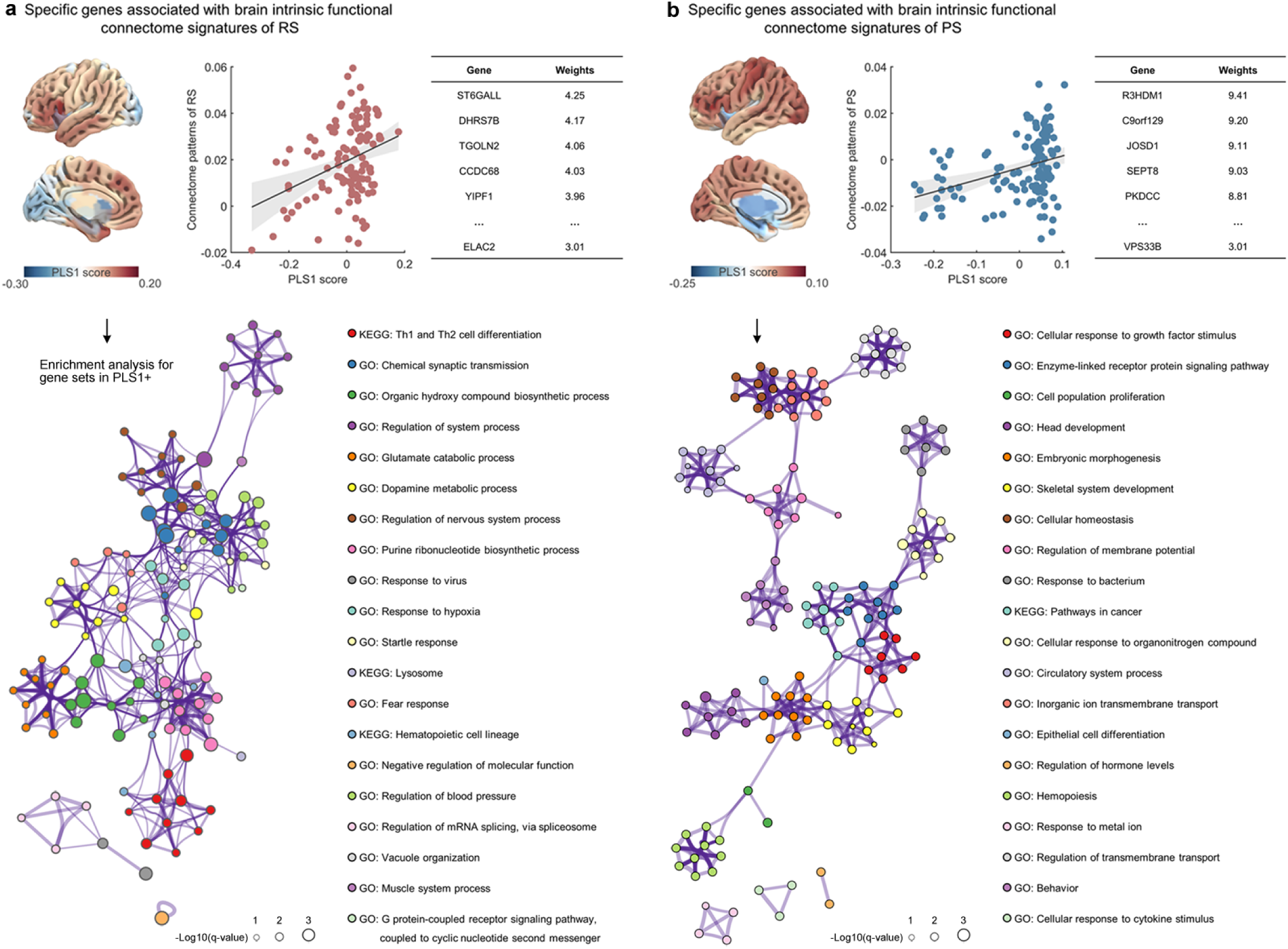
Gene sets related to brain functional connectome patterns of RS and PS were enriched for specific biological processes. **(a)** Regional functional connectome map of reward sensitivity is correlated with the weighted gene expression map of PLS1+ gene set. The genes in the PLS+ gene set are significantly (P_FDR_<0.05, Benjamini–Hochberg correction) enriched for the synapse transmission and metabolic processes, such as “chemical synaptic transmission”, “glutamate catabolic process” and “dopamine metabolic process”. **(b)** Regional functional connectome map of punishment sensitivity is correlated with weighted gene expression map of PLS1+ gene set. The genes in the PLS+ gene set are significantly (P_FDR_<0.05, Benjamini–Hochberg correction) enriched for immune response and stress adaption processes, including “enzyme-linked receptor protein signaling pathway”, “cell population proliferation” and “cellular homeostasis”. The enrichment network in both panel a and panel b represent the similarities within each cluster and between clusters (circle nodes represent enriched biological processes; their size depends on the proportion of genes related to the enriched biological process among all input genes; nodes with the same color belong to the same cluster; nodes within the same cluster are typically similar to each other; biological processes with similarity >0.3 are connected through edges).

The PLS1+ gene set is mainly enriched for the synapse transmission and metabolic processes, such as “chemical synaptic transmission”, “glutamate catabolic process” and “dopamine metabolic process”, whereas the PLS1- gene set is mainly enriched in the processes for maintaining celluar organization and homeostasis such as “homotypic cell-cell adhesion” and “regulation of small molecular metabolic process” (**Fig. S1a**).

In addition, a significant spatial correlation was also observed between the PLS1 (explained variance, 37.85%) and the brain intrinsic functional connectome patterns of PS (r_(121)_=0.34, P_spin_=0.03, **Fig. 3b**), with high gene expressions in the fronto-insular regions that display highly similar rs-FC patterns to indicate PS. After applying the FDR correction to the normalized weights of PLS1, we found 1999 positive weighted genes (Z>3.01) and 2046 negative weighted genes (Z<-3.01, P_FDR_<0.005). The PLS1+ gene set is predominantly enriched in biological processes related to immune response and stress adaption, including “enzyme-linked receptor protein signaling pathway”, “cell population proliferation” and “cellular homeostasis”. In contrast, the PLS1- gene set is primarily associated with inflammation processes, such as “cellular response to cytokine stimulus”, “cell activation” and “inflammatory response” (**Fig. S1b**).

### Relative contributions of specific neurotransmitters to RS- and PS-related brain functional connectome alterations

To further decode the distinct brain functional connectome patterns of RS and PS by chemoarchitectures, we investigated the relationship between neurotransmitter systems and the brain functional connectome patterns of RS and PS using two separate multiple linear regression models. Consistent with prior works ^33,70–72^, eight receptors from five different neurotransmitter systems which play functional roles in the processing of reward and punishment were included as predicted variables in the regression model, including the dopamine (DA1, DA2), serotonin (5HT1a, 5HT1b, 5HT2a), glutamate (mGluR5 and NMDAR), gamma-aminobutyric acid (GABA) and Opioid (MOR, **Fig. 4a**). We found that the model explained 4.18% of the variance in the brain connectome variations of RS. To be specific, several receptors from serotonin (5HT1a, weight=-0.02, p<0.001, 5HT1b, weight=-0.11, p<0.001), dopamine (D2, weight=-0.11, p<0.001), glutamate (weight=-0.03, p<0.001) and GABA (weight =- 0.09, p<0.001, **Fig. 4b**) systems contributed relatively identical and negatively predict the variations of brain connectome of RS. However the 5HT2a (weight=0.23, p<0.001) occupied more relative contribution and interacted with D1 receptors (weight=0.11, p<0.001), as well as the opioid system (weight=0.09, p<0.001, **Fig. 4b**) to positively predict the brain connectome patterns of RS.

**Fig. 4.**
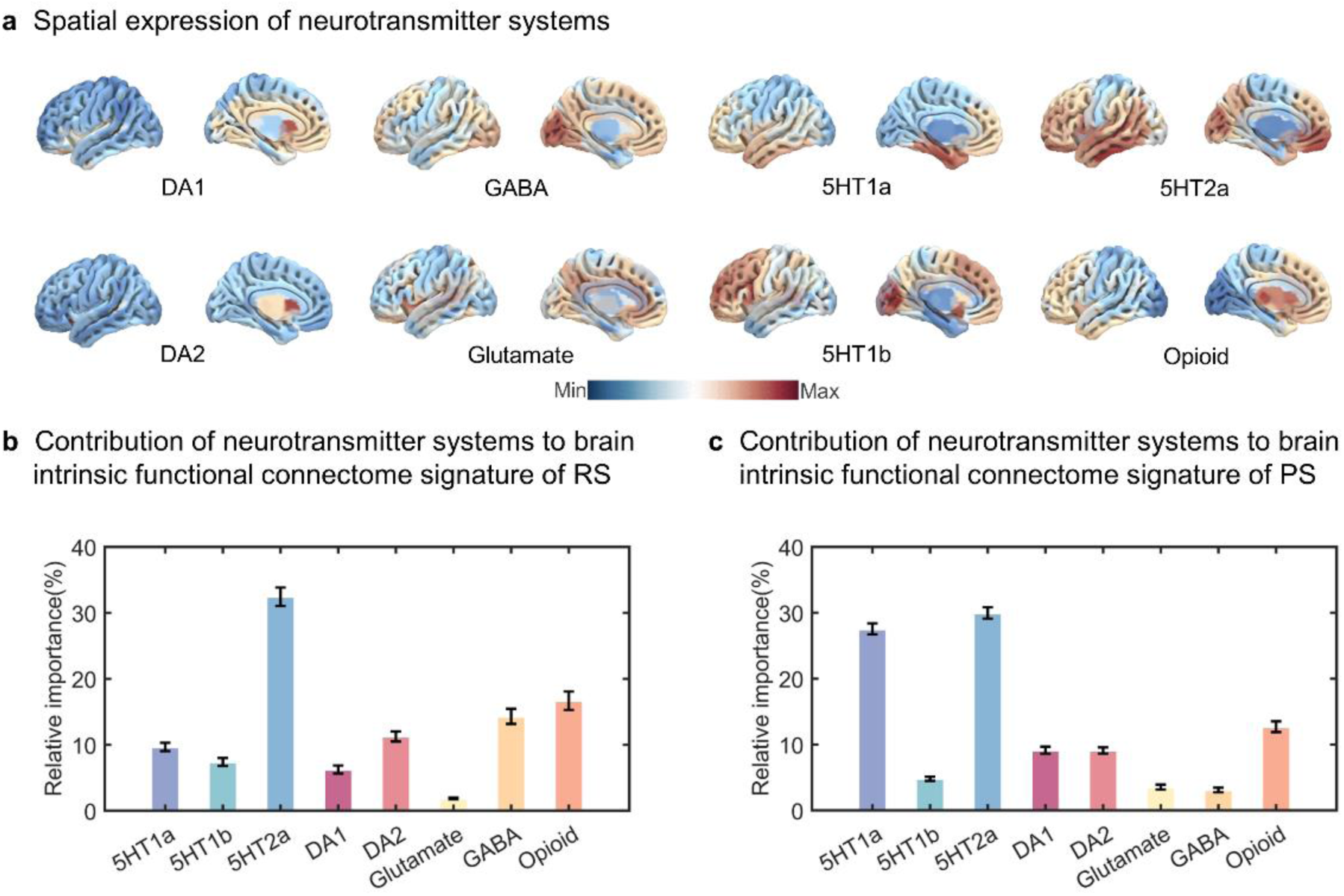
Relative contributions of specific neurotransmitters to brain functional connectome alterations of RS and PS. **(a)** Brain anatomical distribution of different neurotransmitters, **(b-c)** The relative contribution of each neurotransmitter to predict the regional functional connectome changes for reward and punishment sensitivities. The error bars represent standard deviation of bootstrapping-based (5000 iterations) relative importance.

Additionally the second regression model explained 7.04% of the variance in the brain connectome patterns of PS. Within this model, five receptors from serotonin (5HT1a, weight=- 0.27, p<0.001, 5HT1b, weight=-0.03, p<0.001), dopamine (D1, weight=-0.07, p<0.001, D2, weight=-0.03, p<0.001) and opioid (weight =-0.06, p<0.001, **Fig. 4c**) systems negatively predict the variations of brain connectome of PS, while the 5HT1a had more relative contribution.

Moreover, the 5HT2a receptor (weight=0.31, p<0.001) also showed more relative contribution and combined with the neurotransmitter systems of GABA (weight=0.02, p<0.001) and glutamate (weight=0.05, p<0.001, **Fig. 4c**) to positively predict the brain connectome patterns of PS. Taken together, the 5HT2a may serve as a pivotal hub modulating the neural intrinsic representations of RS and PS, with this process critically dependent on its intricate interplay with distinct neurotransmitter systems.

### Replication in an independent sample

To support the robustness of results in the discovery study, we repeated the analyses in the replication sample. Similarly, we found that functional connectome variation of RS and PS were distinctly mapped in the brain, with frontal-striatal network (**Fig. 5a**) being sensitive to track RS change and the IFG being more implicated in PS (**Fig. 5b**). Those functional connectome signatures of RS and PS were also specific to predict the social or monetary reward or punishment-induced activations (**Fig. S2a-S2c**), and further linked with genes that were similarly founded in the discovery sample (RS, odds ratio=20.64, p<0.001, **Fig. 5a**; PS, odds ratio=5.20, p<0.001, **Fig. 5b**). Finally the serotonin potentially acted functional roles together with the opioid, dopaminergic and GABAergic systems to have effects on the functional connectome patterns changes for RS (5HT2a, weight=0.03, p<0.001; DA1, weight=0.12, p<0.001; Opioid, weight=0.11, p<0.001) and PS (5HT2a, weight=0.03, p<0.001; GABA, weight=0.01, p=0.001).

**Fig. 5.**
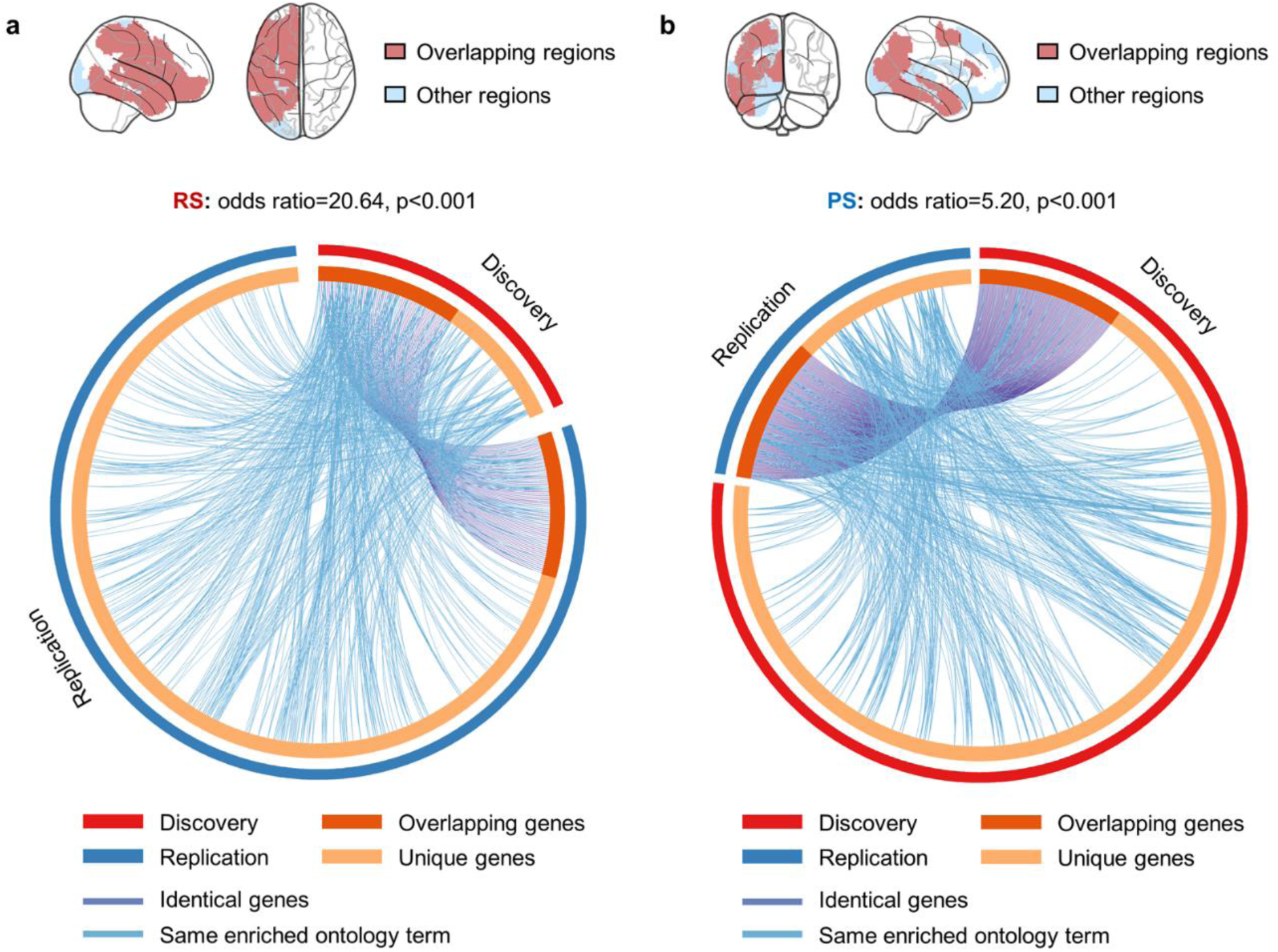
Validating RS- and PS-related brain intrinsic functional connectome patterns, and examining their associations with transcriptional profiles. **(a)** Both discovery and replication samples display similar regional functional connectome maps for reward sensitivity. The genes associated with variations in the functional connectome for reward sensitivity in both discovery and replication samples show a high degree of overlap. **(b)** Both discovery and replication samples display relatively similar regional functional connectome maps for punishment sensitivity. The genes associated with variations in the functional connectome for punishment sensitivity in both discovery and replication samples are highly overlapped.

## DISCUSSION

The hyper-and hypo-reward or punishment sensitivities are strong predictors of motivational and emotional dysfunction related symptoms, having garnered considerable attention as prominent transdiagnostic features of psychopathology ^73,74^. However the lack of an overarching neurobiological characterizations currently limits the early identifications and neuromodulations of abnormal reward and punishment sensitivities. As such we utilized the multimodal fMRI and AHBA microarray datasets in combination with mainstream imaging-transcriptomic analyses to determine the macroscale brain functional connectome signatures of RS and PS, further validate their specificity in social or monetary motivation contexts, and characterize their associations with specific molecular genes and neurotransmitters. By leveraging IS-RSA analysis, we identified dissociable brain functional connectome signatures for RS and PS, with rs-FC patterns in the fronto-striatal network encoding reward responsiveness, while the fronto-insular system was more involved in sensitivity to punishments. The brain intrinsic functional connectome signatures of RS and PS were also specific to differentiate social and monetary reward or punishment events. Applying PLS models to the brain intrinsic functional connectome patterns of RS or PS, and gene expressions maps revealed that variations in rs-FC for RS and PS were linked to the topography of specific gene sets enriched in ontological pathways, including synaptic transmission, dopaminergic metabolism, immune response and stress adaptation.

While on the neurotransmitter level, the 5HT2a as an essential component of serotonin system was a pivotal hub modulating the neural intrinsic connectome representations of RS and PS, with this process critically dependent on its interactions with dopaminergic, opioid and GABAergic systems. Final replication analyses in the independent sample also supported the robustness of the results obtained in the discovery study. Taken together, these findings offer the first comprehensive neural connectome mapping of reward and punishment sensitivities, highlighting their functional engagement in social and monetary reinforcement contexts and linking them to specific molecular mechanisms.

To capture neurofunctional variations associated with RS and PS separately we employed the IS-RSA model on rs-FC patterns and observed dissociable mapping of RS and PS in terms of intrinsic functional connectomes. Specifically the fronto-striatal network was implicated in encoding reward responsiveness, while the fronto-insular system was more strongly associated with sensitivity to punishment. The multivariate IS-RSA conducted based on rs-FC pattern in fact reflects similar temporal associations between spatially distant neurophysiological events across subjects, and has been widely applied to examine functional relationship between neural regions and cognitive traits ^60,61,64^. In this vein, the increased similar coupling within the fronto-striatal circuit potentially reveal that regions within this network are more synchronized in their spontaneous activity to predict reward sensitivity. Supporting that, strong functional connectivity within the fronto-striatal circuitry was positively associated with greater reward sensitivity even in the absence of external task demands ^75^. Furthermore the fronto-striatal network also plays a crucial role in reinforcement-guided decisions, with increased coupling between regions such as the striatum and vmPFC facilitating rewarding decisions that are influenced by reward value evaluation ^76^. In these findings the dorsal and ventral parts of striatum, often considered as the brain’s reward hub, works in concert with the vmPFC to assess the potential value of rewards ^77^ and select actions that have previously led to rewards ^78^, separately. However, in the context of punishment learning, the involvement of the striatum is more complex, with its dorsal part connecting to extensive brain regions to support tasks focused on avoidance learning ^15^, behavioral inhibition and response to negative feedback ^79,80^. While disengaging from behaviors that led to punishment rely on the functions of dorsal striatum, how to initially detect the stimuli that is emotionally aversive via automatic interoceptive awareness ^81^ and how to maintain the consistent avoidance behaviors from punishments or threats critically require the involvement of fronto-insular circuitry. Within this circuitry, the insula plays a key role in the evaluation of aversive states ^82^ such as unexpected monetary loss experience ^83^ and negative feelings like vicarious pain and disgust ^18,84^, whereas the IFG represents higher-order nuclei that relay and integrate an entire array of important signals to resolve conflicts ^85^ and inhibit impulsive actions when individuals receive punishments ^86^. Connections of the IFG with limbic brain structures in particular: 1) drive the punishment decisions for a social normal violation ^87,88^, and 2) reduce behavioral interference of aversive pictures ^89^. Our finding is consistent with these observations and further supports the specificity of fronto-insular circuitry in decoding social or monetary punishment information in independent samples.

Given that convergent evidence outlined the modulatory roles of genetic variation in the processing of reward and punishment stimuli ^90,91^, we modelled the spatiotemporal overlap of neufunctional alterations relevant to RS or PS, and weighted gene expression patterns across the whole brain. Our results indicated that the intrinsic functional connectome variations for RS were associated with gene expression enriched in chemical synaptic transmission and dopamine metabolic processes, whereas the PS-related changes in functional connectome were linked to gene expression enriched in immune response and stress adaption processes. The chemical synaptic transmission refers to the process by which neurons communicate with each other through the release and reception of chemical signals, such as neurotransmitters ^92^. During the situation of reward learning, synaptic transmission is primarily mediated by neurotransmitters such as dopamine which is mainly released in basal ganglia ^93^. It has been suggested that dopaminergic modulation of the direct striatonigral pathway in the basal ganglia are critical in reward learning, while the reversible neurotransmission blockade of this pathway impaired the learning ability of distinguishing associative rewarding stimuli from nonassociative ones ^94^.

Moreover as indicated by one recent excitatory synapse hypothesis of depression, impaired excitatory synaptic transmission particularly reduced activity in the mesolimbic reward circuitry and developed symptoms of depression such as anhedonia and aberrant reward-associated perception ^95^. In contrast the immune response serving as an extensive defense mechanism against harmful threats is particularly sophisticated in protecting individuals from environmental punishments and stressors. Numerous human experimental works have systematically examined the regulatory roles of diverse immune challenges in sensitivity to various punishment stimuli, such as: 1) acute immune responses triggered by lipopolysaccharide rapidly increased the encoding of punishment learning signals and enhanced avoidance of high probability punishment stimuli ^96^, 2) upregulation of inflammation via typhoid vaccination prompted a significant enhancement of insula encoding of punishment predictive errors ^97^, and 3) acute stress induced prolonged inflammatory responses and significantly improved choice accuracy performance on loss trials over the ensuing four months ^98^.

In addition to neurotransmitter level, we observed complex yet relatively concordant associations: the serotonin receptor subtype 5HT2a made important contributions to RS-related functional connectome variations in conjunction with dopaminergic and opioid systems, while distinctly functioning with GABAergic system to predict changes in PS-related functional connectome patterns. Both 5HT and dopamine have long been implicated in learning which behaviors are associated with reward ^99,100^, yet the precise mechanism of how dopamine interacts with serotonin to support rewarding choices remains unspecific. A recent innovative research demonstrates for the first time exactly that dopamine and serotonin work together in opposite ways such that dopamine initiates “go” signal to drive reward-seeking when outcomes exceed expectations, while serotonin acts as a brake (“wait” signal), promoting patience and consideration of long-term benefits ^101^. Both the ‘go’ signal from dopamine and the ‘wait’ signal from serotonin are critically required for human to properly evaluate and grasp rewarding opportunities. Moreover our findings align with previous works that suggested endogenous opioid signaling as the underpinning of hedonic aspects of reward ^102^ by revealing the involvement of opioid neuromodulator in neural functional connectome map related to reward sensitivities. With respect to punishment, early discoveries from animal models provided preliminary evidence for the synergistic functions of serotonin and GABAergic neurons in negative events ^103^ such that: 1) aversive stimuli such as quinine and footshock caused an overall mild inhibition of serotonin neurons but strongly activated GABA neurons ^45^, 2) optogenetic inhibition of GABA cells disinhibited serotonin cells and prevented the acquisition of social avoidance in mice exposed to social threats ^104^. Together these findings underscore the engagement of specific neural signaling systems in tracking neurofuntional alterations related to reward and punishment sensitivities which may represent biological targets for modulating reward/punishment deficits domains.

While prevailing neural models converge on implicating the activations within fronto-striatal and fronto-insular regions in encoding the responsiveness to reward and punishment, separately, we updated these observations by providing a more comprehensive dissection of the neurotranscriptomic dissociation between RS and PS. Specifically we demonstrated that the macroscopic network functional alterations of RS and PS are supported by distinguishable microscale biological pathways. However there are several limitations needed to be considered. First, while we repeated the same analyses in replication sample to support the robustness of initial findings in discovery study and consequently observed relatively consistent findings some differences were also remained. Second, the AHBA dataset utilized in this study provides a limited sample size and includes data only for the right hemisphere for two subjects. Although recent imaging transcriptomic studies indicate that this data is sufficiently generalizable in healthy cohorts ^50,57^, a large sample of whole brain gene expression data is required for future studies. Third, while we characterized the relative contribution of neurotransmitter distributions to predict RS-or PS-related functional connectome patterns, this only indicate the correlational but not the causational information. Despite that recent evidence demonstrated a closely alignment of neurotransmitter profiles with brain structural and functional connectivity patterns ^105^, future studies may test the causation via record and manipulate the activity of multiple neuromodulators to check their functions in contexts of reward and punishment processing ^106^.

To conclude, we provide the first neurobiological characterizations for reward and punishment sensitivities by mapping distinct neural functional connectome patterns within the brain and identifying its linkage with distinguishable transcriptomic profiles. The fronto-striatal network was identified as central to reward processing, while the fronto-insular system was more involved in punishment sensitivity. These patterns were distinctively linked to gene sets enriched in biological process including synaptic transmission, dopaminergic metabolism, immune response, and stress adaptation. Interactions of serotonin with dopaminergic, opioid and GABAergic systems may play complex regulatory roles in shaping neural functional organizations in response to rewards and punishments. Together, this study offers an integrative, multi-scale understanding of behavioral tendencies toward reward and punishment reinforcers, which may inform future early identifications and treatments for psychopathological conditions involving reward and punishment processing deficits.

## METHODS

### Participants and Questionnaires

We utilized a randomized parallel resting state fMRI design on independent discovery and replication samples. The discovery sample included N=450 healthy participants, whose behavioral and neuroimaging datasets were collected between September and November 2019 as part of the Behavioral Brain Research Project of Chinese Personality (BBP) ^107^. All participants were free from a current or a history of psychiatric, neurological, or other medical disorders, had no current or regular use of psychotropic substances including nicotine, had a normal body mass index (18-24.9), had no visual or motor impairments and contraindications for MRI. A total of 23 participants were excluded due to excessive head movement (a mean framewise displacement [FD] larger than 0.3 mm), leading to a final sample of N=427 (287 Female, Mean±SD, age=18.83±0.93 years; 140 Male, age=18.98±1.39 years) included into main analyses.

With the purpose of replication, N=240 healthy participants finished data collection during 2020 and were included based on the same enrollment criteria. However 12 participants were excluded due to excessive head movement, thus the data of 228 participants (155 Female, age= 22.35±2.69 years; 73 Male, age=22.86±1.22 years) were incorporated into the replication analyses.

Regarding the behavioral measures, all participants completed the 48-item Sensitivity to Punishment and Sensitivity to Reward Questionnaire (SPSRQ) which is a self-report scale designed to assess an individual’s sensitivity to reward and punishment in various contexts ^108^. This questionnaire consists of two distinct subscales: one for sensitivity to reward (e.g., how likely a person is to seek pleasurable experiences or avoid missing out on rewards) and one for sensitivity to punishment (e.g., how much a person is motivated to avoid negative outcomes or discomfort). By leveraging the SPSRQ, numerous studies have quantified the RS and PS in large populations ^109^, and specified their effects on multiple cognitive processes such as risk-taking and decision making ^110^.

This study was approved by the Research Ethics Committee of the Southwest University (H24218) and was adhered to the latest revision of the Declaration of Helsinki. All participants recruited for this study were compensated with 50 Chinese Yuan and signed informed consent after being fully briefed about the study’s purpose, benefits, and potential risks.

### Multi-neuroimaging data acquisition and preprocessing

The resting state fMRI (rs-fMRI) dataset in both discovery and replication studies were obtained on a 3.0 Tesla Prisma Siemens scanner located at the Southwest University Center for Brain imaging, whereas the task BOLD fMRI dataset were acquired on a 3.0 Tesla Prisma Siemens scanner at the Temple University Brain Research & Imaging Center ^111^ and on a 3.0 Tesla MAGNETOM Prisma Siemens scanner at the Laboratory of Brain Imaging, Neurobiology Center at the Nencki Institute of Experimental Biology Polish Academy of Sciences ^112^, separately. In line with our previous studies ^107,113^, all rs-fMRI data were preprocessed using the validated workflows in a publicly available CONN functional connectivity toolbox (version 20.b; https://www.nitrc.org/projects/conn) and in SPM12 (Welcome Department of Cognitive Neurology, London, UK; http://www.fil.ion.ucl.ac.uk/spm). For more details about imaging data acquisition and preprocessing, please see **Supplementary Information**.

### Identification of brain functional connectome signatures of RS and PS

Given that individuals exhibit varying responses to reward or punishment stimulus ^2,62^, how to identify the specific neurofunctional alterations of RS and PS while controlling for potential individual differences is a central question. Recent advances in imaging-based methodologies offer promising solutions by recommending the utilization of a novel inter-subject representational similarity analysis. This approach can enable to determine whether different individuals share similar neural representations for a given cognitive process and has been proved robustness in revealing individual variations in affective experiences and social decisions ^60,61,64^. As such we employed the inter-subject representational similarity analysis to examine which brain regions display similar rs-FC patterns in response to the RS or PS. Additionally, considering that the AHBA included only two samples in both hemispheres (left hemisphere only, n=4), our subsequent analyses focused exclusively on the left hemisphere to ensure the robust estimations. We firstly extracted the regional averaged time course from each of the left brain regions defined by the Brainnetome atlas ^114^(105 cortical and 18 subcortical regions) and computed the Pearson correlation coefficients between time courses of each pair of regions.

These correlation coefficients were further normalized using Fisher’s r-to-z transformation, resulting in a 123 × 123 symmetric rs-FC matrix for each participant. Next we constructed a neural representational dissimilarity matrix for each region by calculating the Euclidean distance between each pair of participants based on the rs-FC matrix. Additionally we also created behavioral dissimilarity matrices using the Euclidean distance between each pair of participants in the RS and PS score space, respectively. Finally we estimated the correlation between each regional dissimilarity matrix ad the behavioral dissimilarity matrix of RS or PS using the Spearman’s rank-order correlations on the upper triangle of the matrices. The resulted correlations were considered statistically significant if the p-value was less than 0.05, after applying the Benjamini-Hochberg correction ^115^ for multiple comparisons. Significant correlations were taken to indicate a significant relationship between behavioral distance and regional representation distance, as well as a significant association of RS or PS with intrinsic functional connectivity patterns in a given region.

### Specificity analyses of brain functional connectome signatures of RS and PS in reward or punishment events

To examine whether the brain intrinsic functional connectome signatures of RS and PS could be extended into specific monetary or social reward and punishment contexts, we applied the brain functional connectome patterns of RS and PS to two independent fMRI dataset. In the first dataset (N=52), participants were asked to finish two tasks including: (1) giving responses to visual cues indicating monetary outcomes: high gain, low gain, high loss, low loss, and neutral (no gain or loss) (S1: N=52), and (2) choosing the faces of peers who may potentially like them (social reward and punishment – peers’ like and dislike, S3: 6 participant from S1 were removed due to imaging data loss, resulting in a final sample size of N=46); while in the second dataset, participants only respond to visual cues representing large and small gain or loss (S2: N=37 male). We then modeled those pre-processed BOLD time series datasets using three general linear models (GLMs), each incorporating different regressors tailored for different analytical purposes. The first GLM model that included the presentation onsets of outcome separated by the gain, neutral and loss conditions, as well as nuisance regressors pertaining to the head motion (three rotation parameters and three translation parameters), was established to obtain the contrast images which were used for test whether the brain intrinsic functional connectome signatures of RS and PS can predict individual real-time responses to monetary reward and punishment from neutral events. Similarly, we included regressors for gain and loss outcomes in the second GLM model to assess whether the brain intrinsic functional connectome signatures of RS and PS can differentiate reward and punishment events, or vice versa. While focusing on checking the prediction of brain intrinsic functional connectome signatures of RS and PS in social reward and punishments-related situations, we incorporated regressors corresponding to the positive and negative peer feedbacks conditions, alongside a nuisance regressor for head motion in the third GLM model.

Next, three separate support vector machine classifiers implemented in the Spider toolbox (http://people.kyb.tuebingen.mpg.de/spider) were trained on single subject first-level GLM contrast images to differentiate: (1) between responses to the monetary gain (or loss) and responses to the neutral outcomes, (2) between responses to the monetary gain (or loss) and responses to the loss (or gain) outcomes, and (3) between responses to the social like (or dislike) and responses to the dislike (or like) feedbacks, based on the brain intrinsic functional connectome signature of RS or PS. In order to reduce the chance of overfitting, we employed a leave-one-subject-out cross-validation procedure in which the N-1 subjects were used for training and the remaining subject was used for testing, and we calculated forced-choice classification accuracies between monetary/social reward and punishment outcomes based on the pattern responses within the brain intrinsic functional connectome signatures of RS or PS.

### Association of gene expression with brain functional connectome changes of RS and PS

We capitalized on the full dataset of protein coding genes (n=20,737) from six donor brains as provided by the AHB atlas (http://human.brain-map.org/) to investigate the relationship between brain intrinsic functional connectome patterns of RS and PS, and the transcriptomic files. The microarray expression data were initially processed and mapped to 246 regions from Brainnetome atlas ^114^ according to a reproducible workflows ^116^ via the use of a well-established abagen toolbox ^117^. Next, a regional expression matrix was generated for each donor, with 246 rows representing brain regions and 15,528 columns representing the retained genes.

Consistent with previous works ^56,57^, we averaged the retained genes across all donors and further extracted the gene expression data of left hemisphere (123 x 15,528 matrix) for the following analyses to guarantee robust estimations.

In order to determine the relationship between brain intrinsic functional connectome changes of RS and PS, and the transcriptional activity for all 15,528 genes, we employed the PLS regression models in which the 123 x 15,528 regional gene expression matrix was included as predictor variable, whereas the brain intrinsic functional connectome changes of RS and PS were included as response variables, separately. The first component of PLS (PLS1) was strongly correlated with brain intrinsic functional connectome patterns of RS and PS across 123 regions. To reduce the confounding effects of the spatial autocorrelation, we employed permutation testing based on spherical rotations of the brain intrinsic functional connectome signatures of RS and PS to test the association of PLS1 and RS-or PS-related neurofunctional connectome patterns ^68,69^. A bootstrapping method (5000 iterations) was finally used to estimate each gene’s weighting coefficient in the PLS analysis, and the z-scores for each gene’s weight on PLS1 were calculated by dividing the weight by its bootstrap standard error.

### Enrichment analyses

To identify the functional pathways that are enriched in the PLS1 gene sets that are related to brain intrinsic functional connectome patterns of RS or PS, we conducted gene enrichment analysis using Metascape (https://metascape.org/), an online platform that integrates a variety of bioinformatics tools and enables gene-meta analysis across over 40 independent knowledge bases ^118^. Specifically the genes with weights greater than 3.01 or less than-3.01 (Z > 3.01 or Z <-3.01, All P_FDR_<0.005) were input into the Metascape website for Kyoto Encyclopedia of Genes and Genomes (KEGG) and Gene Ontology (GO) biological process analyses. Metascape then identified ontology terms that contained a statistically greater number of genes in common with the input list than expected by chance using the widely adopted hypergeometric test. A Kappa-test score was further computed as the pairwise similarities between two enriched terms and the resulting similarity matrix was hierarchically clustered to organize the terms into distinct clusters. Finally, the obtained enrichment pathways were thresholded for significance at 5%, corrected by the FDR.

### Association of neurotransmitter systems with brain functional connectome changes of RS and PS

We finally investigate whether the brain intrinsic functional connectome changes of RS and PS were spatially correlated with the several neurotransmitter systems particularly engaged in the reward and punishment processing. In this regard, we utilized whole-brain volumetric PET receptor images from a cohort of over 1200 healthy participants to create spatial maps of 19 distinct neurotransmitter receptor across nine different neurotransmitters ^105^. Based on previous works ^33,70–72^, eight receptors from five different neurotransmitter systems which play functional roles in the processing of reward and punishment were analyzed, including the dopamine (DA1, DA2), serotonin (5HT1a, 5HT1b, 5HT2a), glutamate (mGluR5 and NMDAR), gamma-aminobutyric acid (GABAA) and Opioid (MOR). In accordance with the procedures in Hansen et al., receptors and transporters with more than one mean image of the same tracer (i.e., 5-HT1b and D2) were averaged together in a manner that weights each image by the number of participants in the cohort ^105^ and then generated a neurotransmitter density map.

The eight neurotransmitter density maps were further assigned to 123 regions, and the averaged density value of all voxels within the region was further z-normalized and defined as the region’s neurotransmitter density value. After that we explored the contributions of different neurotransmitters to the brain intrinsic functional connectome changes of RS and PS, respectively, by using a multivariate linear regression model as implemented in R package relaimpo (relative importance of regressor in linear models, version 2.2-5). To address linear regression with multiple collinear regression, we calculated the relative importance metrics and further assessed the relative contribution of each neurotransmitter using a bootstrapping method with 5000 samples. The multiple comparisons were corrected by FDR with a significance threshold of P<0.05.

### Replication analyses

To validate the results obtained in the discovery study, we repeated the analyses in terms of brain intrinsic functional connectome identification for RS and PS, their specificity in social and monetary reward or punishment contexts, and their functional associations with transcriptional profiles in the replication sample.

## Acknowledgements

This work was supported by the National Natural Science Foundation of China (32271123), National Key Research and Development Program of China (2022YFC2705201), the Fundamental Research Funds for the Central Universities (SWU2009104) and Innovation Research 2035 Pilot Plan of Southwest University (SWUPilotPlan006). We sincerely thank all colleagues for their unwavering support during data acquisition and for their invaluable discussions, which greatly contributed to improve the manuscript. We also extend our gratitude to all volunteers who participated in our study, to the Allen Institute for Brain Science for providing the gene expression data and to the OpenNeuro platform for offering the neuroimaging data.

## Author contributions

This study was designed by TX and TF. All data were collected by TX, CZ, XZ and RZ with the help of HC, QH, XL and JQ; TX performed data analysis and results integration with frequent discussions with ZC, FZ, WZ, XG, XC and XZ. The manuscript was initially written by TX and TF and critically revised by all authors.

## Competing interests

All authors declare no competing interest.

## Notes

### Competing Interest Statement

The authors have declared no competing interest.

