## Supplemental Information for "Dissociable neurofunctional and molecular characterizations of reward and punishment sensitivity"

### Supplemental Methods

#### Multi-neuroimaging data acquisition

The resting state fMRI (rs-fMRI) dataset in both discovery and replication studies were obtained on a 3.0 Tesla Prisma Siemens scanner located at the Southwest University Center for Brain imaging, whereas the task BOLD fMRI dataset were acquired on a 3.0 Tesla Prisma Siemens scanner at the Temple University Brain Research & Imaging Center [1] and on a 3.0 Tesla MAGNETOM Prisma Siemens scanner at the Laboratory of Brain Imaging, Neurobiology Center at the Nencki Institute of Experimental Biology Polish Academy of Sciences [2], separately. In line with our previous studies [3,4], all rs-fMRI data were preprocessed using the validated workflows in a publicly available CONN functional connectivity toolbox (version 20.b; https://www.nitrc.org/projects/conn) and in SPM12 (Wellcome Department of Cognitive Neurology, London, UK; http://www.fil.ion.ucl.ac.uk/spm).

*Rs-fMRI and T1-weighted structural MRI scan*: To characterize intersubject similarity of rs-FC patterns, each subject from discovery and replication samples underwent an approximately 8 min of rs-fMRI scan (240 volumes) using a blood oxygen level-dependent (BOLD-weighted) sequence: repetition time (TR)/echo time (TE)=2000 ms/30 ms, flip angle=90°, field of view (FOV)=224 × 224 mm$^2$, resolution matrix=112 × 112, slice thickness=2 mm, voxel size=2 × 2 × 2 mm$^3$, slices=62, phase-encoding direction=PC » AC. To facilitate alignment of individual subject images into a common space, a high-resolution T1-weighted structural image was obtained using a three-dimensional gradient sequence: TR/ TE=2530 ms/2.98 ms, flip angle=7°, FOV=256 × 256 mm$^2$, resolution matrix=256 × 256, slice thickness=1 mm, voxel size=0.5 × 0.5 × 1 mm$^3$, slice oversampling=33.3%, slices=192, phase-encoding direction=AC » PC. During the rs-fMRI scanning, subjects were instructed not to think of anything special, maintain fixation on the crosshair displayed on the screen, and stay in a fixed position.

*Task BOLD fMRI and T1-weighted structural MRI scan*: In order to examine the prediction of brain connectome signatures of RS and PS to monetary and social reward or punishment events, we included two validated cohorts who performed the monetary incentive delay and social door tasks, respectively. For the first validation cohort (N=52, https://openneuro.org/datasets/ds004920/versions/1.1.1), the high-resolution structural scan was acquired (sagittal plane) with a T1-weighted magnetization prepared rapid acquisition gradient echo (MPRAGE) sequence: TR/TE=2400 ms/2.17 ms, voxel size=1.0 × 1.0 × 1.0 mm$^3$, flip angle=8°, FOV=224 x 224 mm$^2$, while the functional T2*-weighted images were acquired using a simultaneous multislice (multi-band factor=2) gradient EPI sequence: TR/TE=1750 ms/29 ms, voxel size=3 × 3 × 3 mm$^3$. In addition, a B0 fieldmap was collected to unwarp and undistort functional images (TR=645 ms; TE1=4.92 ms; TE2=7.38 ms; matrix=74 × 74; voxel size=2.97 × 2.97 × 2.80 mm$^3$; 58 slices, with 15% gap; flip angle=60°). For the second validation cohort (N=37 males, https://openneuro.org/datasets/ds005479/versions/1.0.2), the T1-weighted anatomical scans were performed using Echo Planar Imaging sequence: TR/TE=2300 ms/2.96 ms, voxel size=1x1x1 mm$^3$, whereas the task T2*-weighted scan was performed using Simultaneous multislice imaging sequence (397 volumes): TR/TE=2000 ms/30 ms, slice thickness=2.5 mm, 60 slices acquired in interleaved order, Multiband acceleration factor=2.

**Multi-neuroimaging data preprocessing**

*Preprocessing of resting-state BOLD fMRI data*: All functional images in discovery and replication study were initially processed through following steps: (1) slice timing correction, (2) motion correction and susceptibility artifact correction based on field map, (3) realignment of 3D anatomical data to Montreal Neurological Institute (MNI) space via the Diffeomorphic Anatomical Registration Through Exponentiated Lie Algebra method, and (4) spatial smoothing with a 6 mm full-width at half-maximum Gaussian kernel. The remained functional images were further denoised using the anatomical component-based correction method [5]. Specifically, signals from cerebrospinal fluid and white matter (five principal components), and movement parameters (six motion parameters, six temporal derivatives, and their squares), as well as the linear trend were regressed out as confounds [6]. Data scrubbing was then performed to avoid head motion concerns, whereas the bad time points were also flagged based on a criteria of any volumes with FD>0.5 mm as well as the two succeeding volumes and one preceding volume to reduce the spillover effect of head motion [6]. Finally, the resulting functional images were applied for a temporal band-pass filter (0.008–0.09 Hz) to minimize the effects of very low-frequency drifts and high-frequency noises, followed by the linear detrending.

*Preprocessing of task BOLD fMRI data*: The task functional images were preprocessed and analyzed using standard protocols in SPM 12. The first 5 volumes of each functional time series were discarded to allow for T1 equilibration. Remaining images were corrected for acquisition time delay, realigned to correct for head motion, unwarped for magnetic field inhomogeneities correction, and co-registered with the T1-weighted structural image. After that the images were normalized to MNI standard space (interpolated to 2 × 2 × 2 $mm^3$ voxel size) using the segmentation parameters from the anatomical images. In line with previous studies on successfully decoding cognitive and affective processes [7-9] via the use of multivariate pattern analyses, the normalized images were spatially smoothed using an isotropic Gaussian kernel with full-width at half-maximum of 8 mm. This smoothing process has been demonstrated to improve inter-subject functional alignment while preserving the sensitivity to mesoscopic activity patterns that are consistent across individuals [10,11].

### Supplemental Results

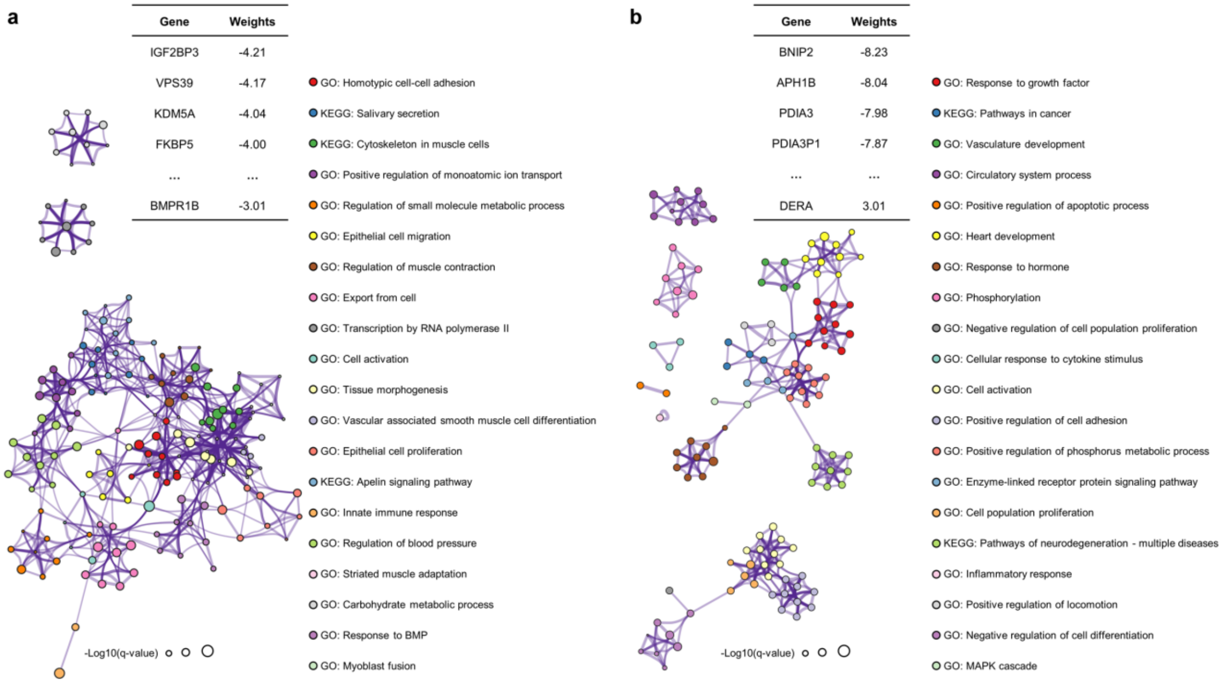


**Fig. S1 Functional enrichment analyses for PLS1- gene sets in replication sample. (a)** Regional functional connectome map of reward sensitivity is correlated with weighted gene expression map of PLS1- gene set. The genes in the PLS- gene set are significantly ($P_{FDR}$<0.05, Benjamini–Hochberg correction) enriched in the processes for maintaining celluar organization and homeostasis such as "homotypic cell-cell adhesion" and "regulation of small molecular metabolic process". **(b)** Regional functional connectome map of punishment sensitivity is correlated with weighted gene expression map of PLS1- gene set. The genes in the PLS- gene set are significantly ($P_{FDR}$<0.05, Benjamini–Hochberg correction) enriched for inflammation processes, such as "cellular response to cytokine stimulus", "cell activation" and "inflammatory response". The enrichment network in both panel a and panel b represent the similarities within each cluster and between clusters (circle nodes represent enriched biological processes; their size depends on the proportion of genes related to the enriched biological process among all input genes; nodes with the same color belong to the same cluster; nodes within the same cluster are typically similar to each other; biological processes with similarity >0.3 are connected through edges).

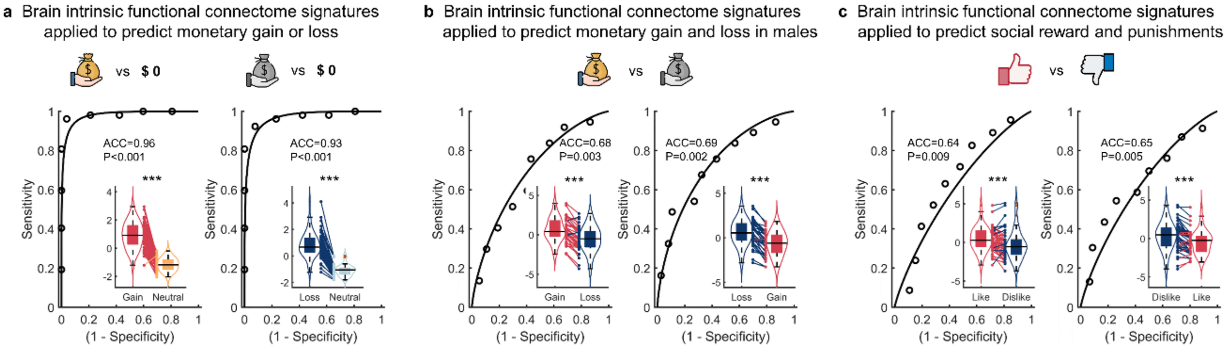


**Fig. S2 Validating the specificity of RS- and PS-related brain functional connectome signatures within social/monetary reward and punishment contexts in replication sample. (a-c)** Robust specificity of the brain functional connectome signatures of reward and punishment sensitivities in the contexts of social and monetary reward or punishments, separately. The violin and box plots show the distributions of multivariate neural pattern map response to classify multiple events (i.e., Monetary gain vs neutral, Monetary loss vs neutral; Monetary gain vs loss, Monetary loss vs gain; Social like vs dislike, Social dislike vs like). The box is bounded by the first and third quartiles, and the whiskers stretched to the greatest and lowest values within the median±1.5 interquartile range, while each colored line between dots represents each participant's paired. ***p<0.001
